# Delta and Theta Activity Provide Shared and Unique Sensitivity to Alcohol and Cocaine Use Disorder

**DOI:** 10.64898/2026.06.04.723639

**Authors:** Devin Butler, Edward Bernat, Nadia Mattanah, Sophia Nahabedian, Vaughn Steele

**Affiliations:** University of Iowa; University of Maryland, College Park; Yale University

## Abstract

The P300 amplitude reduction, or P3AR, is a decrease in the magnitude of the P300 event-related potential (ERP) waveform, typically measured via EEG during oddball tasks, and widely observed in those diagnosed with Alcohol use Disorder (AUD). Though widely acknowledged as a biological correlate of AUD, most of the extant literature on the P3AR features conventional time-domain measurement approaches, and further, the work has predominantly focused on alcohol use specifically. In order to precisely capture the nature of this association as well as assess generalizability to other substances, the current study utilized advanced time-frequency analyses to compare P3 activity in individuals with AUD, cocaine use disorder, and comorbid alcohol and cocaine use. Time-frequency analyses indexed separable theta and delta activity underlying the conventional time-domain P3. Findings revealed that in the conventional time-domain, we observed P3AR shared across alcohol and cocaine use. These findings replicate literature suggesting P3AR associated with substance use. Within time-frequency analyses, we found broad delta reductions across substances, aligning with literature suggesting that delta is mostly comprised of the P3 ERP. For theta-band activity, we found reductions in. amplitude specific to alcohol users only, with no significant diminutions present for cocaine users. As most previous literature focused on alcohol use, unique theta modulations relative to cocaine users provides theoretical clarity in terms of the association of P3AR to substance use more broadly. The findings of the current study highlight the utility of time-frequency measurement approaches in parsing overlapping activity and indexing separable processes with relevance to substance use otherwise unaccounted for in the time domain.

## Introduction

A widely observed biological correlate of alcohol use disorders is a reduction in the amplitude in the P300 (P3) event-related potential (ERP) component, referred to as the P3 amplitude reduction (P3AR) (Alper, 1999; Anokhin et al., 2000; Bauer, 1997; Bauer & Hesselbrock, 1999; Begleiter et al., 1984; Iacono et al., 2002; Kempel et al., 2003; Porjesz, Begleiter, & Garozzo, 1980; Singh et al., 2009). However, with a majority of the work on P3AR focused on alcohol use and its effect on P3 amplitude (Cohen et al., 1995; Hamidovic & Wang, 2019; Porjesz, Begleiter, & Garozzo, 1980; Singh & Basu, 2009), ERP studies on substances other than alcohol are relatively few in number (Singh and Basu, 2009), especially in regards to studies on the effects of cocaine use on P3. Though studies do exist examining the effect of cocaine use on the P3 that show decrements in a similar fashion to alcohol (Bauer, 2001; Biggins et al., 1997), there are discrepancies, as mixed results and inconclusive findings remain, limiting definitively conclusive statements. For example, some studies report on substance use broadly, as opposed to singular cocaine use (Wan et al., 2010); others report results showing no P3 reduction for cocaine users, even showing a P3 increase (Spronk et al., 2016); or reporting no P3/ERP analyses (Franken et al., 2008; Pike et al., 2015). Thus, of the few studies that exist in exploring this relationship to cocaine, discrepancies stemming from inconsistent sampling, methodology, and results have kept the exact etiology and significance of these findings unclear. Importantly, the majority of P3AR studies assess the relationship using conventional time-domain component measurement approaches (Keil et al., 2022), while time-frequency (TF) approaches can index separable theta and delta activity underlying conventional component measures (Bachman & Bernat, 2018; Bernat et al., 2007, 2015). As the P3 in an oddball task is largely made up of separable theta and delta activity, in the present study, we directly compared P300 activity, using TF analyses, among individuals characterized by either cocaine, alcohol, or comorbid cocaine and alcohol use disorder, relative to healthy controls with no drug or alcohol dependence in order to evaluate relative effects of each on the P3. We analyzed in the conventional time-domain as well as delta and theta-frequency bands. We highly expect to replicate well-studied diminutions in theta and delta for alcohol use (Porjesz & Begleiter, 1995), while filling in a critical gap in the literature regarding our understanding of P3AR relative to cocaine use.

### Broad Overview of Alcohol and Cocaine: Why They Require Independent Assessment with P3AR

A large number of studies have appeared in the literature documenting the effects of alcohol use on the P3 ERP (Cohen et al., 1995; Porjesz, Begleiter, & Samuelly, 1980; Porjesz & Begleiter, 1995), such that subjects with alcohol use disorder have reduced P3 amplitudes. Though this P3 amplitude reduction (P3AR) has been associated extensively with alcohol use, studies on the P3 effects of cocaine use are relatively few in number (Bauer, 2001; Singh & Basu, 2009). Singh and Basu (2009) note that the P300 has strong potential as an endophenotype for substance use disorder (SUD), but it has only deeply been studied in relation to alcohol specifically, as opposed to other substances. Although previous literature has identified reductions in P3 amplitude in individuals with substance use and broad externalizing problems (Bernat et al., 2020; Gao & Raine, 2009; Hajcak et al., 2010; Patrick et al., 2006; Singh & Basu, 2009), independent evaluation of alcohol and cocaine use is necessary given their differential effects on body and cognition. This study evaluates alcohol and cocaine separately to assess their associations with the P3AR. Clarity in how the two substances variably affect individuals and measuring ERPs across users of these specific substances may reveal more about the cognitive processes indexed by the P3 response and its amplitude reduction among externalizing disorders.

Cocaine and alcohol are among the most frequently used substances in the United States as well as globally – 1.34 billion people consumed harmful amounts of alcohol worldwide in 2020, and 21.5 million people used cocaine at least once in the same year in the United States (*IHME, 2022*; *UNODC World Drug Report 2022 - World | ReliefWeb*, 2022). Importantly, concurrent use of alcohol and cocaine is popular among drug users (Pennings et al., 2002). The physiological impact of both alcohol and cocaine is well understood: Both alcohol and cocaine can cause changes in the mesolimbic dopaminergic system which can affect reinforcement of addictive behaviors (Mash et al., 2002). These drugs also influence the presence of dopamine in parts of the brain activated while interpreting rewarding stimuli (Di Chiara, 1997). This drug-induced reward system activation may encourage a person to repeat the behaviors that caused the positive sensations in the brain (Di Chiara, 1997). Repeated activation in turn elicits likings and cravings that manifest into addictive behaviors, such as repeated use and dependence on alcohol and cocaine (Sarafino & Smith, 2022). Cocaine and alcohol are both frequently used addictive substances that alter chemical pathways in the brain.

Alcohol and cocaine do have differences, however, in terms of psychological and physiological effects on individuals, as they oppositely affect the reward pathways in the brain through their impact on dopamine (Ma & Zhu, 2014). Alcohol, a nervous system depressant, causes disinhibition of dopamine release in the nucleus accumbens, which can affect one’s alertness, judgment, emotions, and motor skills (Ciucă Anghel et al., 2023). Conversely, cocaine, a nervous system stimulant, accelerates and amplifies neural processes by blocking the reuptake of dopamine in the brain (Mash et al., 2002; Ciucă Anghel et al., 2023). Through increased extracellular dopamine, the drug elicits euphoric sensations and increased alertness (Ciucă Anghel et al., 2023). The divergent impact of these two commonly used substances on dopamine pathways might suggest the presence of separable underlying neural mechanisms. The current study seeks to investigate if separable cognitive processes are engaged in cocaine and alcohol users, as reflected in EEG P3 activity.

As alcohol has been the most commonly studied in terms of association with P3AR, we address a literature gap here by also assessing cocaine use, and it represents a shift that should continue onto addressing the P3 relationship to opiate use. As the opioid crisis has been declared a public health emergency (HHS, 2022), with 1.6 million people having an opioid use disorder as recently as 2020, it is incredibly useful to explore how different substances relate to P3. Further, as evidence suggests P3 decrements are related not just to substance use, but broader externalizing (Patrick et al., 2006) and general psychopathology (Bernat et al., 2020), how different substances relate to psychopathology is a crucial avenue of research to understand the relationship substance use has with cognitive and affective processing and other psychopathology.

### P3AR: A Broad Overview

P3AR has been observed broadly (Porjesz et al., 2005; Polich et al., 1994). This includes alcohol use, as noted above, but also relative to psychopathology more broadly, including psychotic thinking (Turetsky et al., 2000), schizophrenia (Jeon & Polich, 2001, 2003), externalizing problems (Iacono et al., 2003), depression (Sara et al., 1994), and anxiety with more mixed findings (Bauer et al., 2001; Kimble et al., 2000). Recent work has related P3AR to emerging models of psychopathology, notably the hierarchical taxonomy of psychopathology (HiTOP) (Kotov et al., 2017). Earlier work in this area demonstrated relationships with a broad externalizing factor, including alcohol use, antisocial behavior, and disinhibitory problems (Nelson et al., 2011; Patrick et al., 2006). More recent work from our group has suggested that P3AR may be better understood as indexing the shared variance across all psychopathology (cf. *p*-factor; Bernat et al, 2020). As we have seen the P3AR to alcohol use, and to externalizing broadly, this effect suggests a system is present driving P3 down, and it is critical to study other substances to examine their role and the extent to which it is similar or different from the current literature. As this recent work helps us more deeply understand the P3AR relationship with both alcohol use and psychopathology, the current study seeks to understand how different substances, such as cocaine, relate to this relationship. As the P3 is broadly related to psychopathology, one might expect that the same relationship persists for cocaine use, but it has been insufficiently studied, and the current study attempts to explore this possibility.

P3AR has been widely studied in the context of alcohol use, with many findings suggesting it is a good phenotypic marker (Begleiter et al., 1984; Hesselbrock et al., 2001; Polich et al., 1994), but it has only been studied as an endophenotype in alcohol use disorders and not other substances. With evidence suggesting that P3 activity is an objective indicator of brain activity disrupted by acute effects of cocaine use (Bauer, 2001; Kouri et al., 1996), further investigation is necessary into other substances to understand how the P3AR operates more broadly. The focus of P3AR and substance use has focused exclusively on alcohol, and in this study, we wanted to contrast the effects of cocaine and alcohol use on P3 amplitude. As the two substances are commonly used together, this dataset features groups of independent cocaine and alcohol users, as well as polyusers that meet criteria for alcohol and cocaine use disorder.

### P3AR and Substance Use: Research Largely Focused Alcohol, Less for Cocaine and Other Substances

ERP studies largely revolve around alcoholism, with less focus on other substances (Singh and Basu, 2009). Decades of research have continued to affirm the association of P3AR with alcoholism (Porjesz et al., 1980; Cohen et al., 1995; Hamidovic and Wang, 2019), with research showing the P300 deficit in numerous circumstances, including both after abstinence and in the sons of individuals with alcohol use disorder even if they were naïve to alcohol and other drug use. Meta-analyses and replications of these findings have continued to confirm these consistencies and this association of P3AR to alcohol use, identifying the P300 as a phenotypic marker for alcohol use (Begleiter et al., 1984; Fairbairn et al., 2021; Hesselbrock et al., 2001; Porjesz, Begleiter, & Samuelly, 1980; Rangaswamy & Porjesz, 2014; Singh & Basu, 2009). Overall, alcohol’s effects on P3 amplitude is highly researched and well-validated, with consistent research going towards identification of mechanisms underlying alcohol use.

In contrast, there is much less research on ERPs and cocaine use, especially regarding the P300 ERP. For the relatively few studies involving cocaine, inconsistencies in methodology and results remain. One study examined the late positive potential, or LPP (indexing incentive salience and motivational significance of stimuli) and found an amplitude increase in response to cocaine stimuli (Franken et al., 2008); another study reported decreased inhibitory control in cocaine users relative to healthy controls, but this was not an ERP study (Pike et al., 2015); Kouri and colleagues (1996) found no differences in P3 activity in substance using (cocaine and heroin) patients and healthy controls when presented for detoxification, but did find P3AR for patients afterwards. Alper noted that cocaine use in humans is most commonly associated with increased beta frequency band activity (Alper, 1999). Wan and colleagues found reduced P3 amplitude for substance users, but this study examined the P300’s ability to predict treatment completion, and did not focus on any specific substance, while including alcohol users in the sample (2010). Another study reported that cocaine did not affect parietal P3 amplitude, while actually showing an increased prefrontal P3 (Spronk et al., 2016). One study examined patients with histories of cocaine and alcohol dependence, and found no difference in P3 between cocaine and cocaine and alcohol-dependent subjects, but did find impaired P3 in the two patient groups relative to controls, only to Novel stimuli, with no Target analyses reported (Biggins et al., 1997). These findings show that differences in both methodology (e.g., limited samples of only dually-dependent users, comorbid psychiatric disorders, family histories of alcoholism, treatment completing patients) and results highlight a gap in knowledge for the P300 relationship with cocaine use. Overall, research has found a significant correlation between a reduced P3 response and alcohol use (Begleiter et al., 1984; Iacono et al., 2002; Porjesz, Begleiter, & Garozzo, 1980), but the relationship between P3AR and cocaine use is more limited and inconclusive.

Bauer (2001) compared P300 activity among individuals characterized by histories of cocaine, or cocaine and alcohol, opioid dependence or no previous drug or alcohol dependence; they found P3AR in all patient groups relative to healthy controls. This study’s results suggest that cocaine effects on the P3 operate similarly to alcohol’s well-studied association. This finding combined with the inconsistent and few cocaine ERP studies noted above serve as motivation for the current study, where a major goal was to examine the extent of how similarly or contrastingly cocaine use might affect P3 relative to alcohol. With these substances being of two different drug classes, with differential physiological effects, and the literature having conclusive evidence on the relationship between alcohol use and P3 while Bauer (Bauer, 2001, 2002) suggesting similar decrements in cocaine with a backdrop of inconsistent reporting, we sought to explore any shared or unique sensitivities from the use of these two substances. Very few studies have examined the effects of cocaine on the P300 measures of cognitive functioning, and with the effects varying, a study that explores the effects of a stimulant such as cocaine on the P3 is warranted.

### Time-Frequency Analyses of the P300

The P300, or P3, is a widely-studied event-related potential (ERP) in basic and clinical studies (Bashore & van der Molen, 1991), and it is observed as a positive-going amplitude maximally occurring around 300 milliseconds (ms) following stimuli presentation (Polich, 2007). The P3, most extensively studied in “oddball” paradigms in which participants attend to target stimuli that are presented infrequently among other non-target and/or standard stimuli (Hajcak et al., 2010), is often used as a measure of cognitive resource allocation and activation of cognitive schemas to process task information and evaluate stimuli, ultimately indexing salience detection, attention allocation, information processing and working memory (Bernat et al., 2020; Brennan & Baskin-Sommers, 2018; Polich, 2007).

In order to precisely contrast effects of alcohol and cocaine on the P3, time-frequency analyses must be utilized. Most research on the P3 (broadly, as well as the P3AR) has been done with time-domain methodology, where components are defined by the amplitude occurring within specific temporal start and end points (Bachman and Bernat, 2018). This approach faces difficulty in dissociating temporally overlapping ERP activity, as these measures may feature energy from earlier ERPs that continue during the time of the P3. Discrepancies in time-domain feedback components can be clarified with time-frequency (TF; an important tool for assessing electrical brain activity from event-related paradigms) analytic approaches, thus using only time-domain measures may be insufficient in many contexts. TF analyses can characterize activity simultaneously in time and frequency, providing measures that are separated in frequency, but which overlap in time. The gap in literature regarding cocaine use and P3AR can be addressed through TF measures.

Numerous studies have shown the P3 in oddball tasks can be understood as a mix of early frontal theta band activity (around 3 to 7 Hz) and later posterior delta band activity (around 0 to 3 Hz) (Bachman & Bernat, 2018; Başar-Eroglu et al., 1992, 2001; Bernat et al., 2007; Demiralp et al., 2001; Gilmore et al., 2010; Jones et al., 2006). Both theta and delta have been related to oddball responses. The overall processing sequence involves an anterior theta response first, more closely tied to orienting, and then a posterior delta response more closely tied to cognitive processing. The P3 has largely been researched in the time domain, but time-frequency measures showed it is comprised of separable theta and delta activity, allowing for a deeper understanding of processing.

To explore how similarly cocaine’s effects are on the P3 relative to alcohol, we must use time-frequency analyses, as the P3 can be more optimally represented by multiple overlapping TF components (Gilmore et al., 2010). Gilmore and colleagues’ findings suggest TF representations of EEG/ERP may produce more optimal indices of underlying neurophysiological processes. As noted above, delta and theta parsing has shown that delta and theta provide different roles and indices in stimuli processing. P3-related delta, which tends to be parietally maximal and makes up most of the P300 (Bachmann and Bernat, 2018), has been considered to index signal matching, decision-making, and memory updating (Basar-Eroglu et al., 1992; Karakas, Erzengin, & Basar, 2000). We have particular interest in the theta frequency band, as our group has presented findings that show differential effects between theta and delta when applying TF analyses, with recent evidence from our lab suggesting that both the P3 and theta are independently related to general psychopathology (Milandu et al., 2023). Theta in the P3 window has been considered to index focused attention and memory encoding processes (Başar-Eroglu et al., 1992; Klimesch, 1999; Yordanova et al., 2000). Theta is more closely associated with orienting and salience processing, and these functions may be implicated in regard to psychopathology. With cocaine classified as a stimulant that impacts an individual’s awareness and attentional allocation through perceived hypervigilance, theta response may be unique when compared to alcohol users’ theta response, seeing as alcohol is nervous system suppressant.

Time-frequency analyses have shown alcohol use is related to reductions in both theta and delta amplitude (Jones et al., 2006; Rangaswamy et al., 2007). In a review by Rangaswamy and Porjesz (2014), it is conclusively stated that similarly to observations from above-mentioned time-domain P3 studies, there too exists theta and delta reductions related to a broad range of alcohol problems such as alcoholism, binge drinking, long-term abstinence, as well as adult and adolescent risk. Further, this evidence highlights reduced theta and delta as excellent trait markers for alcoholism, similarly with the P3AR. TF delta and theta analyses provide a comparable level of information to ERP analyses, and these findings have been seen across a number of paradigms, including oddball (Andrew & Fein, 2010). Porjesz and colleagues reviewed that P3 amplitude is largely comprised of theta and delta oscillations (Porjesz et al., 2005). Further, they reviewed that alcohol use is associated with significantly reduced theta and delta amplitudes, respectively. As TF theta and delta frequencies are related to different cognitive operations (delta – decision-making, memory updating; theta – encoding, orienting, salience), research examining separable and/or overlapping effects is vital in any detailed examination of consequences of substance use more broadly in cognitive processing. As this much is understood in relation to alcohol, a major aim of the current study was to assess if processes were similarly attenuated with cocaine use, which is much less studied in this manner (Singh and Basu, 2009).

In our current understanding of the P300 ERP, the later posterior delta band activity generally encompasses the main part of the familiar P300 waveform that occurs in the time-domain (Jones et al., 2006; Bachman and Bernat, 2018), whereas the earlier theta does not have a time-domain equivalent in terms of waveform presentation. Our understanding of the P3AR in relation to alcohol use may follow an expected pattern in the delta and theta frequency bands, but the relationship of P3AR to cocaine use in TF is less understood. The P300 is made up of theta and delta, and the time-domain P3AR provides insight into our understanding of a delta P3AR in alcohol users, but there still remains a gap in theta’s role in the association between P3 and substance use more broadly.

While the relationship between P3AR and alcohol has been identified, it is less understood how TF analyses can effectively partition different cognitive processes indexed by the P3 in an oddball task. A major goal of the present study was to elucidate the relative effects of substance use on the P300 ERP within different frequency bands. More specifically, our goal was to contrast the effects of cocaine, and alcohol dependence in an auditory oddball task within the delta and theta frequency bands separably.

### Current Study

The aim of the present study is to explore the relative effects of cocaine and alcohol use on the P3. To ensure accurate and complete characterization of the P3, we applied TF analyses to assess differential effects for cocaine and alcohol in separable theta and delta frequency bands, with the specific goal of assessing any differences between cocaine and alcohol users in P3 amplitude, contributing new insights to the literature that has otherwise reported inconsistent findings on cocaine’s effects, as well as an imbalanced proportion of research dedicated to alcohol and cocaine, respectively. Ultimately, the goal is to understand whether alcohol and cocaine have unique processes. We sought to replicate well-established findings regarding alcohol and P3AR, while addressing a critical gap in the literature regarding the association between P3 and cocaine use. P3AR in alcohol users is explained by both theta and delta oscillations that underlie the ERP in an oddball paradigm (Porjesz & Begleiter, 2003), thus we expect to replicate these findings, while assessing the extent to which cocaine use has a similar effect. Positive findings would clarify discrepancies in findings related to cocaine use in relation to P3AR, as well as provide new insights into information processing and deficits in different substance using groups.

## Methods

### Participants

Participants consisted of 154 healthy adults (75 men) ranging in age from 21 to 59 years (*M* = 36.93, *SD* = 10.30) drawn from the Olin Neuropsychiatry Research Center at the Institute of Living Hartford Hospital and the surrounding community of Hartford, CT via advertisements, presentations at local universities, and word-of-mouth. Using the Structured Clinical Interview for the DSM-IV, all participants were free of any history of psychiatric illness (Axis I and II) and reported no history of psychosis in first-degree relatives. Participants reported they all had normal hearing, and visual acuity was normal or corrected to normal, using contact lenses or MR compatible glasses. Of the 154 participants in the final sample, 79 participants were in the Healthy Control (HC) group. Of the remaining participants, 45 were apart of the Cocaine group, as they meet criteria for Cocaine Use Disorder. 53 participants met criteria for Alcohol Use Disorder, thus they were in the Alcohol group. 23 of the Alcohol and Cocaine users met criteria for both respective use disorders, and they are included in a Polysubstance (referred to as Poly) use group. The remaining single-substance cocaine and alcohol users, 22 and 30, respectively, are hereby referred to as Cocaine (or Coc) and Alcohol (or Alc) groups. 59% of the sample self-identified as White, 22% as Black/African American, 3% as Asian, and 15% as Unknown racial heritage. In terms of identified ethnicity, 10% identified as Hispanic, and 13% Unknown, with the remaining 77% identifying as Non-Hispanic.

### Clinical Assessments

Participants were Alcohol or Cocaine Users, or a part of a Healthy Control group that used neither substance. Diagnoses for substance use disorders were based on criteria from the Structured Clinical Interview for DSM-IV-TR, Axis I disorders (SCID I/ P) (First et al., 2016). 79 participants were in the healthy control (HC) group. For the purposes of the current study, the 22 participants that simultaneously met criteria for cocaine use disorder while not meeting criteria for alcohol use disorder are referred to as the Cocaine group. The 30 individuals meeting criteria for alcohol use and not cocaine use disorder are referred to as the Alcohol group. The overlapping 23 substance users that met criteria for both are referred to as the Poly group.

### Oddball Task

Participants completed a three-stimulus auditory oddball task while undergoing EEG data collection. More recent use of the oddball task incorporates a three-stimulus version with strong evidence suggesting that the third category representing infrequent novel stimuli is compelling for contributing to our understanding of P3 reductions related to clinical problems (Polich, 2004), including substance use (Rodriguez Holguin et al., 1999). Data for each participant were gathered during two consecutive runs of the task. Auditory stimuli were delivered via headphones (Philips SHP2500) and consisted of frequent standard low-pitched (1000 Hz) tones (Standards; 80% of trials), infrequent target high-pitched (1500 Hz) tones (Targets; 10% of trials), and infrequent nontarget novel stimuli (Novels; 10% of trials), which were computer-generated sounds (static, bells, beeps, varied in tone). Each stimulus was presented for 250 ms in pseudorandom order with a variable interstimulus interval averaging 2,000 ms. Participants were instructed to respond as quickly as possible to high-pitched target tones by pressing a single button with their right index finger on a response box. They were further instructed to inhibit responses to any of the other stimuli. A short practice session familiarized participants with the high and low tones, and assured task instructions were understood. The task is designed such that general performance is high, even in severe clinical cases. The analyses for this studied featured only Target and Novel conditions.

### Electroencephalography Recording and Reduction

Electrophysiological data were collected using two Windows-compatible computers and a 64-channel BioSemi ActiveTwo amplifier. The first computer used Presentation software to deliver the stimuli, accept responses, and send digital triggers to the other computer indicating when a stimulus or response occurred. The second computer acquired electroencephalographic data using BioSemi software and amplifier. All signals collected with this BioSemi system were low-pass filtered using the standard fifth order sinc filter with a half-power cutoff of 204.8 Hz then digitized at 1024 Hz during data collection. EEG activity was recorded using sintered Ag-AgCl active electrodes placed in accordance with the 10-20 International System (Jasper, H.H., 1958). The participant’s nose was used as the reference. Six electrodes were placed on the participants face to measure electro-oculogram. These electrodes were placed above, below, and lateral to the canthus of each eye. All offsets were kept below 10 kΩ. After placement of the electrodes, participants were seated in a comfortable chair 60 cm away from a computer monitor on which task stimuli were presented, and were instructed to refrain from excessive blinking or moving during data acquisition. Participants then performed the Oddball task described above.

Pre-processing included down sampling to 512 Hz, bad channel detection and replacement, epoching, eye-blink removal, and low-pass filtering at 15 Hz. Bad channels were identified as having activity four standard deviations away from the mean of all other non-ocular channels. These channels were replaced using the mean of surrounding electrodes. This method, used previously (Anderson et al., 2015; Steele et al., 2014, 2015), was implemented to identify very large artifacts and remove them from the data before applying more stringent data cleaning steps in post-processing. ERP epochs were defined in relation to stimuli presentation, from 1000 ms pre- to 2000 ms post-stimulus. The epoched data were eye-blink corrected using an independent component analysis (ICA) technique. The ICA utility in the EEGLab software (Delorme & Makeig, 2004) was used to derive components then, using an in-house template matching algorithm (Jung et al., 2000), blink components were identified and removed from the data.

ERP components were baseline corrected using a -150 ms to -10 ms window. A low-pass filter at 50 Hz was applied to the data before analysis. A time-frequency principal components analysis was conducted on the data (TF-PCA). An unfiltered, 3-component TF-PCA solution was extracted, accounting for 52% of the variance. Activity in medial frontal theta was best characterized in TF-PC1 (tPC1), and centroparietal delta was best characterized in TF-PC2 (dPC2) of the decomposition components (*p*’s < .05). To reduce ambiguity among PCA measures, theta and delta PCs will include either a “t” or “d” preceding the PC number, respectively. Further, theta and delta measures, referenced simply as Theta and Delta, will be referencing tPC1 and dPC2. Mean measurements from 9-electrode clusters surrounding FCz and Cz (for midfrontal theta and centroparietal delta, respectively) were extracted and used for time-frequency PCA (TF-PCA) (Bernat et al., 2005) analysis described below.

### Time Domain Components

Grand average waveforms for target and novel conditions were computed for each of the 64-electrodes across all participants. Using these grand average waveforms, the P3 window was defined as the maximum positive deflection in the ERP waveform occurring between 203 to 453 ms post-stimulus onset (with ms corresponding to bins of 128 Hz re-sampled signal), fitting a window consistent with the P3 peak. Consistent with conventional literature (Comerchero & Polich, 1999; Kiehl et al., 2001; Molnár, 1994; Polich, 2007), mean measurements from the P3 peak across a 9 electrode cluster surrounding CPz was used to capture the parietal P3.

### Time-Frequency Components

The method used to isolate activity during the P300 window in theta and delta was identical to that of Bernat et al. (2011) and Nelson et al. (2011) (Bernat et al., 2011; Nelson et al., 2011). Specifically, the condition-averaged stimulus-locked ERP signals were filtered using a 50 Hz Butterworth filter (all filters implemented with the matlab butter and filtfilt functions, matlab version 7.4, Mathworks, Inc.) as apart of an unfiltered TF-PCA. Not pre-filtering the ERP revealed in the decomposition separable activity in theta and delta frequencies, respectively. In each PC, activity was appropriately isolated to best represent theta and delta-P300. Next, each filtered signal (theta and delta) was transformed into a time-frequency energy distribution (surface) with the binomial reduced interference distribution variant of Cohen’s class of time-frequency transforms (Bernat, et al., 2005). This was done using full epochs of −1 to 2 s, relative to feedback onset, in order to provide sufficient data to resolve frequencies at and around 1 Hz.

For each of these time-frequency bands, principal components analysis was applied to an area corresponding to the 0 to 750 ms time range and 0 to 10 Hz frequency range; this yielded equivalent time windows for decomposition, but with filters having narrowed the frequency activity within the window to either theta or delta, as described above. Principal components analysis (PCA) was used to identify the primary activation component in each frequency band (delta, theta), corresponding to the largest principal component emerging from the principal components analysis (i.e., the component accounting for the greatest proportion of shared covariance across all time-frequency points). Details for this application of PCA to time-frequency surfaces have been previously published (Bernat, et al., 2005). Briefly, this involves first vectorizing the time-frequency surfaces (e.g., concatenating each frequency row end to end) such that the columns of the data matrix index data from different time-frequency points while the rows index the condition averages (separated by subject and electrode). The PCA decomposition is then conducted using this matrix, utilizing the covariance approach and varimax rotation. The vectorized components are then reassembled into time-frequency surface matrices for interpretation.

The variance accounted for by the first principal component (32.59%) did not substantially exceed that accounted for by the next component (13.23%), and so on for the third principal component (6.92%), indicating that a 3-component solution was justifiable in this case. These time-frequency-based theta and delta principal components scores served as the primary dependent measures in the analyses of brain reactivity to feedback stimuli reported below. Principal component scores were calculated as the mean across the principal component weighted time-frequency surface, in the same manner that principal component scores are calculated with questionnaire data. Electrodes FCz and Cz were most proximal topographically to the maximum of the theta and delta target and novel condition differences, respectively, so data from 9-electrode clusters centered around these electrode sites were employed in the statistical analyses of time-frequency component scores reported below.

### Data Analysis

One-way ANOVAs were conducted for Target and Novel trials between 4 groups: an Alcohol group of participants that solely met for Alcohol Use Disorder, a Cocaine group that met solely for Cocaine Use Disorder, a Polysubstance (Poly) group of individuals that met for both Cocaine and Alcohol Use Disorder, and a Healthy Control group of non-substance users. These ANOVAs were conducted to evaluate differences in unfiltered time domain, theta, and delta frequency band activity during the P3 among these groups. Separate wilcoxon rank-sum tests were then conducted to assess differences in each substance using group’s (SUD) ERP activity relative to healthy controls. Finally, linear polynomial contrasts were applied between the substance using groups (Alcohol, Cocaine, and Poly) to assess the relationships between the Poly and single substance using groups.

## Results

### Event-Related Potentials

Separate one-way ANOVAs were conducted to assess for group differences in both conventional time-domain (to capture P3AR) as well as delta and theta-filtered TF-PCA components for Target and Novel stimuli, respectively.

### Time Domain ANOVA: Group x P3 Activity in Targets

In the conventional unfiltered time-domain, there was a significant effect of SUD group on P3 activity in Target trials [*F*(3, 150) = 3.87, *p* = .011, *ηp2* = .072], such that there was a significant P3 amplitude reduction for each respective substance use group (Alc: *M*: 9.48, *SD*: 1.21; Coc: *M*: 9.88, *SD*: 1.43; Poly: *M*: 9.39, *SD*: 1.38) relative to healthy control (HC: *M*: 13.19, *SD*: .745) (at the *p* < .05 level), as each group had significantly lower P3 than HC, respectively. There were no differences in activity between the SUD groups (i.e. Alcohol, Cocaine, and Poly group P3 activity was not significantly different).

### Time Domain ANOVA: Group x P3 Activity in Novels

In the Novels time domain, effects are similar to the Targets [*F*(3, 150) = 4.56, *p* = .004, *ηp2* = .084], albeit slightly weaker for cocaine users relative to HC: there is still a significant effect of group on P3 activity, and each group’s P3 activity (Alc: *M*: 9.79, *SD*: 1.09; Coc: *M*: 11.21, *SD*: 1.27; Poly: *M*: 9.93, *SD*: 1.25) is lower relative to HC P3 activity (HC: *M*: 13.71, *SD*: .673), but at the *p* < .05 level, the difference between the Cocaine and HC group’s activity was marginally significant, at *p* = .08. These results, taken across targets and novels, suggest there is a broad P3AR effect for SUD groups relative to HC.

**Figure 1.**
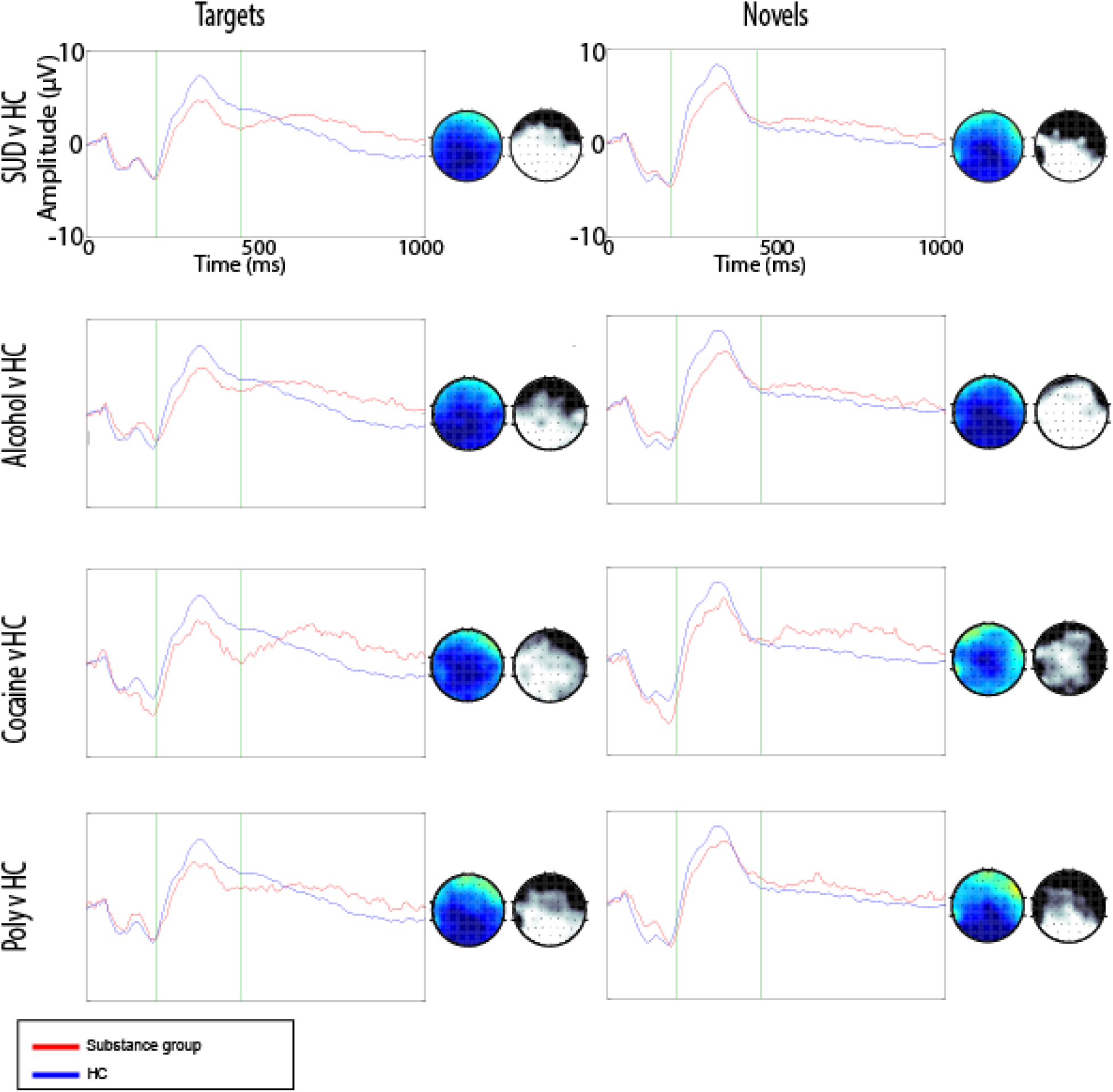
Group comparisons for time-domain decompositions in the P3 ERR Waveforms for both targets (left column) and Novels (right column) comparing activations in a substance use group relative to Control in each row. Magnitude of regional activation is shown by the colored topomaps and statistical significance is shown by the gray scale topomaps.

### Time-Frequency Analysis: Group x Theta Activity in Targets

A one-way between subjects ANOVA was conducted to compare the effect of group on theta activity in Target trials in Alcohol, Cocaine, Poly, and Control/HC groups. There was a significant effect of SUD group on theta activity at the p < .05 level for the 4 conditions [*F*(1,154) = 3.26, *p* = .023, *ηp2* = .061]. Post hoc comparisons using the LSD test indicated that the mean score for the Alcohol group (*M*: 5.19, *SD*: 2.15) was significantly lower than both the Control group (*M*: 11.59, *SD*: 1.33) and Cocaine group (*M*: 14.69, *SD*: 2.51), respectively. However, the Alcohol group did not differ from the Poly group (*M*: 9.31, *SD*: 2.46) significantly. Further, the Cocaine group mean did not differ significantly from the Poly group or the Control group. Taken together, these results suggest a diminution of theta for Alcohol users relative to Control, and there is no diminution of theta for Cocaine users relative to control.

**Figure 2.**
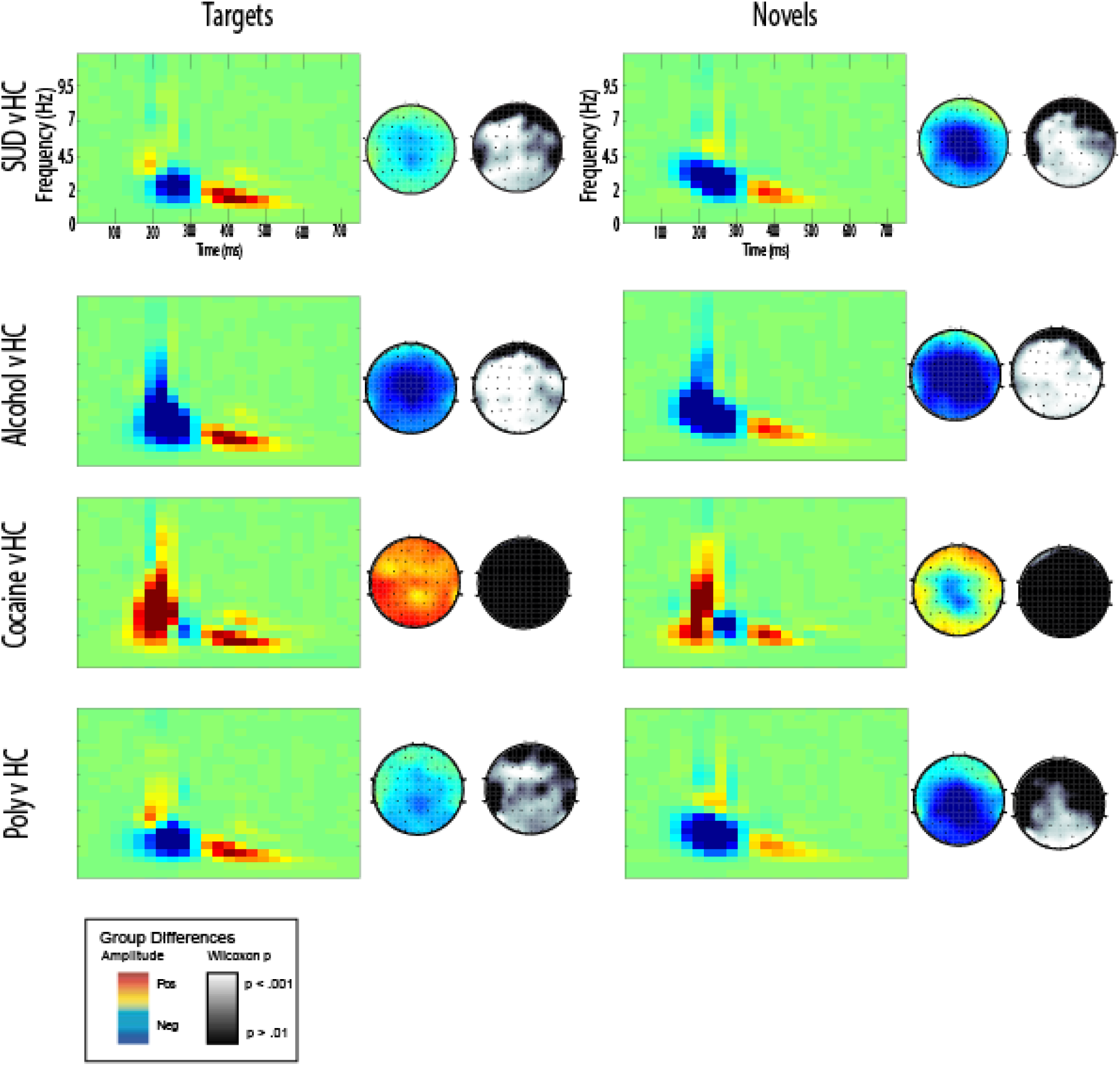
Group comparisons for time-frequency decompositions in the theta band: Topographical surfaces for both targets (left column) and Novels (right column) comparing activations in a substance use group relative to Control in each row. Magnitude of regional activation is shown by the colored topomaps and statistical significance is shown by the gray scale topomaps.

### Time-Frequency Analysis: Group x Theta Activity in Novels

A one-way between subjects ANOVA was conducted to compare the effect of group on theta activity in Novel trials. There was a marginally significant effect of SUD group on theta activity at the p < .05 level for the 4 conditions [*F*(1,154) = 2.33, *p* = .076, *ηp2* = .045]. Post hoc comparisons using the LSD test indicated that the mean score for the Alcohol group was significantly lower than the Control. The Alcohol group (*M*: 12.22, *SD*: 2.86) did not differ from the Cocaine group (*M*: 18.84, *SD*: 3.34), p = .135. Further, the Alcohol group did not differ from the Poly group (*M*: 14.63, *SD*: 3.27) significantly. The Cocaine group did not differ significantly from the Poly group or the Control group (*M*: 20.40, *SD*: 1.76). Taken together, these results suggest a diminution of theta for Alcohol users relative to Control, and there is no diminution of theta for Cocaine users relative to control.

### Time-Frequency Analysis: Group x Delta Activity in Targets

A one-way between subjects ANOVA was conducted to compare the effect of group on delta activity in Target trials. There was a significant effect of SUD group on delta activity at the p < .05 level for the 4 conditions [*F*(1,154) = 4.60, *p* = .004, *ηp2* = .084]. Post hoc comparisons using the LSD test indicated that activity in the Alcohol (*M*: 7.58, *SD*: 2.32) and Poly (*M*: 7.94, *SD*: 2.65) groups was significantly lower relative to the HC group (*M*: 16.09, *SD*: 1.43), respectively. The Cocaine group (*M*: 11.70, *SD*: 2.71) had reduced activity relative to the Control, but this effect was not significant (*p* = .154). The SUD groups did not differ from each other. The results suggest across SUD group, there was a reduction in Delta relative to HC.

### Time-Frequency Analysis: Group x Delta Activity in Novels

A one-way between subjects ANOVA was conducted to compare the effect of group on delta activity in Novel trials. There was a significant effect of SUD group on theta activity at the p < .05 level for the 4 conditions [*F*(3, 150) = 3.98, *p* = .009, *ηp2* = .074]. Post hoc comparisons indicated that the mean score for the Alcohol group (*M*: 7.84, *SD*: 1.69) was significantly lower than the Control group (*M*: 13.45, *SD*: 1.04), and activity did not differ between Alcohol and other SUD groups, respectively. The Poly group (*M*: 7.73, *SD*: 1.93) also differed from the Control group, but the Cocaine group (*M*: 11.71, *SD*: 1.97) did not differ. Taken together, these results suggest a diminution of delta for substance users relative to Control.

**Figure 3.**
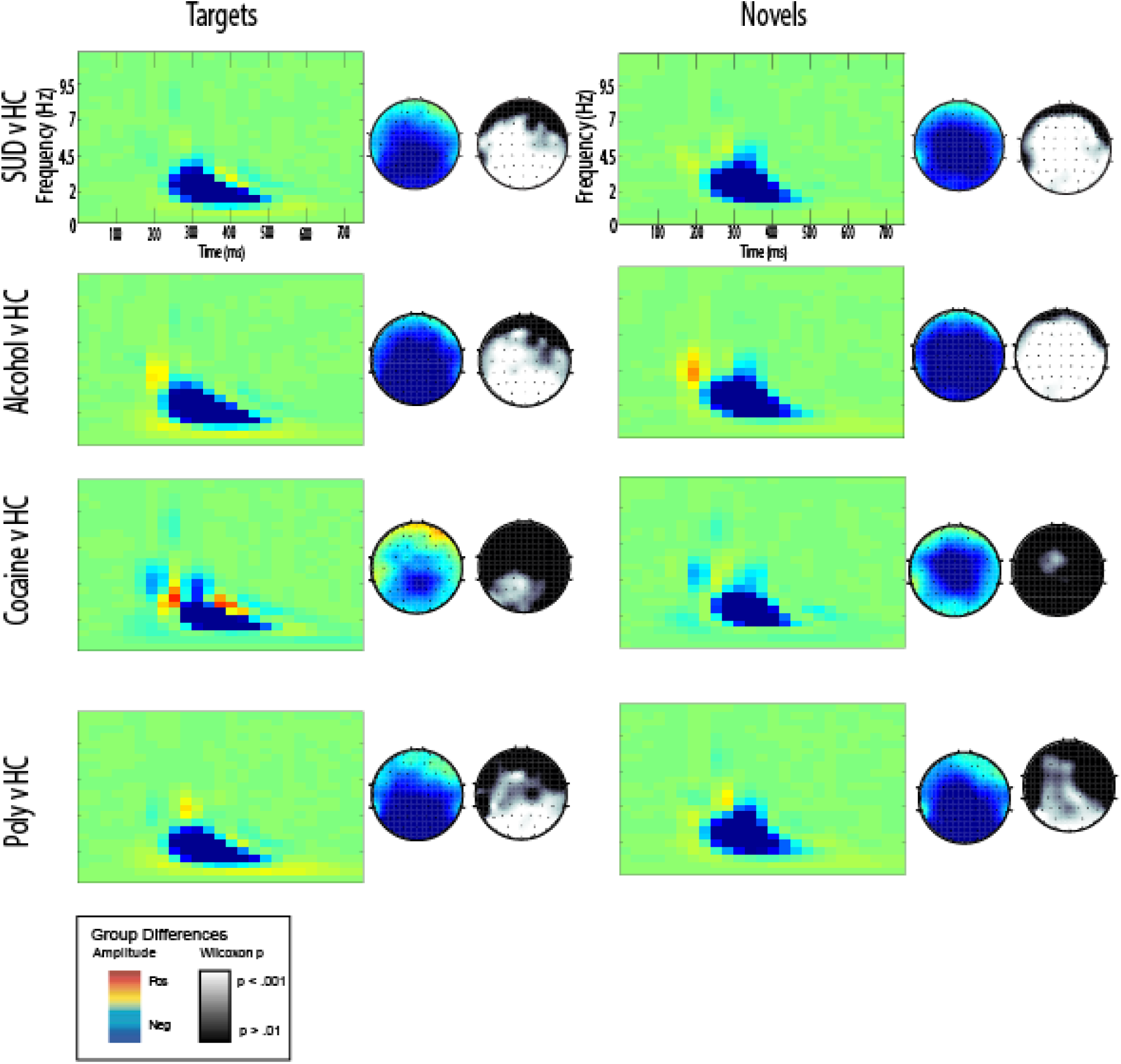
Group comparisons for time-frequency decompositions in the delta band: Topographical surfaces for both targets (left col um n) and Novels (right column) comparing activations in a substance use group relative to Control in each row. Magnitude of regional activation is shown by the colored topomaps and statistical significance is shown by the grayscale topomaps.

### Wilcoxon Analyses: Assessing SUD Activity Relative to HC

Separate wilcoxon rank-sum, or Mann Whitney U tests were conducted to assess for group differences in both conventional time-domain (to capture the P3AR), as well as delta and theta-filtered TF-PCA components for Target and Novel stimuli, respectively. The comparisons in which group differences were calculated are as such: SUD vs HC, Coc vs HC, Alc v HC, Poly vs HC, and Coc v Alc. PCs that primarily represent theta and delta were identified in each decomposition, primarily described by theta-PC1 and delta-PC2.

In the time-domain analysis, a Wilcoxon rank-sum test revealed that SUD amplitude was significantly smaller than HC amplitude in both Targets and Novels. Analyses revealed no amplitude differences between cocaine users and alcohol users.

Similarly, In the delta-filtered time-frequency analysis, SUD amplitude was significantly reduced relative to HC Targets and Novels, save for Cocaine relative to HC in Novels, where the effect was nonsignificant (Table 2).

**Table 1.** Time-Domain P3 Wilcoxons.

|  | Targets |  | Novels |  | Targets and Novels |  |
| --- | --- | --- | --- | --- | --- | --- |
|  | *Z* | *p* | *Z* | *p* | *Z* | *p* |
| **SUD v HC** | -3.27 | 0.001 | -3.29 | < .001 | -3.71 | < .001 |
| **Poly v HC** | -2.27 | 0.023 | -2.25 | .024 | -2.56 | .011 |
| **Coc v HC** | -2.03 | 0.042 | -1.74 | .081 | -2.11 | .034 |
| **Alc v HC** | -2.53 | 0.011 | -2.84 | .005 | -3.05 | .002 |
| **Coc v Alc** | -0.648 | 0.517 | -.741 | .459 | -.019 | .985 |

**Table 2.** Delta Wilcoxons.

|  | Targets |  | Novels |  |
| --- | --- | --- | --- | --- |
|  | *Z* | *p* | *Z* | *p* |
| **SUD v HC** | -3.58 | < .001 | -2.74 | .006 |
| **Poly v HC** | -2.78 | .005 | -2.23 | .026 |
| **Coc v HC** | -1.88 | .061 | -.444 | .657 |
| **Alc v HC** | -2.82 | .005 | -2.89 | .004 |
| **Coc v Alc** | -.407 | .684 | -1.65 | .099 |

**Table 3.** Theta Wilcoxons.

|  | Targets |  | Novels |  |
| --- | --- | --- | --- | --- |
|  | *Z* | *p* | *Z* | *p* |
| **SUD v HC** | -2.34 | .019 | -2.36 | .018 |
| **Poly v HC** | -2.12 | .034 | -1.84 | .066 |
| **Coc v HC** | -.420 | .675 | -.173 | .863 |
| **Alc v HC** | -2.94 | .003 | -2.73 | .006 |
| **Coc v Alc** | -2.41 | .016 | -1.98 | .047 |

In the theta-filtered time-frequency analysis, amplitude was significantly smaller in the Alcohol group and the overall Substance User group relative to healthy control in both Targets and Novels respectively, but there were no differences in amplitude between the Cocaine and HC group. Analyses revealed amplitude was significantly different between cocaine users and alcohol users in both Targets and Novels, in that Cocaine group amplitude was higher than the Alcohol group.

### Linear Polynomial Contrast to Assess Rel. Between SUD Groups

Polynomial contrasts applied to One-way ANOVAs for comparing SUD group (Alcohol, Cocaine, Poly) effects on amplitude were conducted. These contrasts were used to assess group differences in TF measures relative to each other, as opposed to healthy controls. For delta, in both Targets [*F*(2, 74) = 1.29, *p* = .283] and Novels [*F*(2, 74) = 1.92, *p* = .153], there was no difference in amplitude between these groups. Combined with our above one-way ANOVAs, these findings suggest a broad delta reduction for SUD groups. For target trials in Theta [*F*(2, 74) = 3.86, *p* = .026, *ηp2* = .097], we did see a significant effect. Pairwise comparisons revealed that neither Alcohol nor Cocaine group amplitude differed from Poly group (*M*: 9.31, *SD*: 2.46), but Alcohol (*M*: 5.19, *SD*: 2.15) and Cocaine group (*M*: 14.69, *SD*: 2.51) theta target amplitude differed from each other, such that Alcohol had lower theta amplitude. These results confirm a diminution of theta amplitude in Alcohol users relative to Cocaine users, with Poly users nominally in the middle, representing a mixing of the two contrasting effects. In Novel trials, there were no significant group amplitude differences, though approaching trend [*F*(2, 74) = 2.33, *p* = .104]. Pairwise comparisons within the effect show Alcohol (*M*: 12.22, *SD*: 2.86) and Cocaine group (*M*: 18.84, *SD*: 3.34) differences approaching trend in the same direction as the Target effects (*p* = .113), suggesting a similar but weaker Novels effect. Again, at the *p* < .05 level, the Poly group (*M*: 14.63, *SD*: 3.27) showed no significant differences to Alcohol or Cocaine groups but fall nominally within the middle.

## Discussion

### Conventional Time-Domain: P3AR for Alcohol and Cocaine Users

P3AR has been observed broadly, generally within the conventional time-domain. This observation of P3AR is notably associated with not only alcohol and drug use (Polich et al., 1994; Porjesz et al., 2005), as noted above, but also psychopathology more broadly, including psychotic thinking (Turetsky et al., 2000), schizophrenia (Jeon and Polich, 2001, 2003), externalizing problems (Iacono et al., 2003; Patrick et al., 2006), depression (Sara et al. 1994), and anxiety (with more mixed findings) (Bauer et al., 2001; Kimble et al., 2000). Patrick and colleagues have suggested the relationship between the P300 and these facets are entirely attributable to the broad, general externalizing factor (Patrick et al., 2006). Recent work has related P3AR to emerging models of psychopathology, notably the hierarchical taxonomy of psychopathology (HiTOP; Bernat et al., 2020), with work suggesting a P3AR relationship with not only a broad externalizing factor, but also as an index for shared variance across all psychopathology (cf. *p*-factor) (Bernat et al., 2020; Hosch et al., 2023).

Considering this literature, this study’s findings of reduced P3 amplitude in the time domain for both cocaine and alcohol use provides support for the existence of a shared process in the conventional time domain measures. Our findings of P3AR shared across alcohol and cocaine use suggests that this effect is driven by the association with more severe psychopathology (Bernat et al., 2020; Patrick et al., 2006). The current findings are in line with numerous studies showing the P3AR associated with alcohol use (Begleiter et al., 1984; Cohen et al., 2002; Glenn et al., 1996; Patrick et al., 2006; Polich et al., 1994; Porjesz, Begleiter, & Garozzo, 1980), and though fewer in number, for cocaine use as well (Biggins et al., 1997; Bauer, 2001). Reductions in P3 amplitude observed for both cocaine and alcohol users supports the inference of a consistent effect relative to healthy controls, suggesting that though P3AR has been mostly studied in the context of alcohol, a similar process disrupts P3 in alcohol users and cocaine users. These findings are important, as alcohol problems don’t typically occur in isolation (Kessler et al., 1997; Robins & Regier, 1991; Sher & Trull, 1994), and cocaine is another widely-used substance. Combined with the extant literature, the present results support the view that P3AR is not highly specific, and instead appears to relate to a very broad array of problem behaviors and psychopathology, including conduct disorder (Bauer & Hesselbrock, 1999; Patrick et al., 2006), adult antisocial behavior (Patrick et al., 2006), depression (Diner et al., 1985; Singh et al., 2000), and anxiety (Bauer et al., 2001; Xu et al., 2014), as well as latent internalizing and externalizing behaviors representing comorbidities among problem behaviors (Bernat et al., 2020; Patrick et al., 2006). We thus expected to see P3AR across most different kinds of substance use.

### Time-Frequency Analysis: Disentangling Effects in Cocaine and Alcohol

While we found P3AR for both alcohol and cocaine users in the conventional time domain, our findings demonstrate how time-frequency analyses can disentangle nuanced changes in P3 activity. TF research has indicated that energy in centroparietal delta and frontocentral theta can explain the majority of the variance in P3 in target detection/oddball tasks (Porjesz et al., 2005; Rangaswamy et al., 2014; Basar-Eroglu, et al., 2001), and delta and theta index separable processes, providing different functional roles in stimulus processing (Bachman & Bernat, 2018; Bernat et al., 2006; Pandey et al., 2016). Our study represents novel findings that both contribute to and expand upon the current literature: time-frequency analyses precisely disentangled conventional time-domain findings in this study, as, especially within the Target condition, delta findings generally showed the traditional P3AR shared across substances, but the theta-band showed unique activity relative to cocaine and alcohol users. As the P3 is mostly comprised of delta (Bachman & Bernat, 2018; Karakaş et al., 2000; Verleger, 1988), reductions in delta for alcohol and cocaine users serves as validation for the shared process in our time domain findings, in line with the abundance of aforementioned research associating P3AR with substance use. With our delta and time domain findings, this common shared process suggests the P3AR as an overall objective biomarker for substance use, and perhaps externalizing and psychopathology more broadly (Bernat, 2020; Patrick et al., 2006). As for differences in the theta band, findings are potentially related to the unique neurophysiological effects of alcohol and cocaine, respectively. Alcohol, a nervous system depressant, affects people’s judgment, alertness and motor skills; conversely, cocaine is a stimulant that speeds up and amplifies neural processes while eliciting euphoria and amplifying alertness, with cocaine users often reporting that they feel their mental ability actually improves when intoxicated (Siegel, 1984b, 1984a). Cocaine has shown evidence to increase response inhibition and the ability to suppress actions (Fillmore et al., 2005, 2006; Garavan et al., 2008; Spronk et al., 2016).

Neuropsychological impairments associated with alcohol use is well-documented, especially in learning and memory, which are processes indexed by the P3 (Shelton & Parsons, 1987), whereas the nature of cognitive deficits in cocaine users is less understood, with more mixed results (Goldstein et al., 2004). Given that substantial work implicates that theta P3 responses are closely tied to attention allocation, memory updating, orienting, and salience processing (Başar-Eroglu et al., 2001; Bernat et al., 2015; Harper et al., 2014, 2016; Jones et al., 2006; Watts et al., 2017), that we found no theta modulation in cocaine users relative to healthy controls may be reflective of the hypervigilant effect cocaine is understood to produce (Ciucă Anghel et al., 2023; Mash et al., 2002). Alcohol users have depressed judgment and alertness, evidencing an inability to orient in a way that healthy controls can, while cocaine users maintain that vigilance and orienting response (i.e. relative to controls). Taken together, time-frequency analyses revealed that delta was similarly modulated, contributing to P3AR, suggesting both alcohol and cocaine effects not seen in our time domain findings as well as current P3AR time domain literature: alcohol suppresses one’s ability to effectively orient and process stimuli, whereas cocaine users are able to stay more readily alert to allocate attention in the same manner as the healthy controls, even showing nominal increases in theta amplitude relative to healthy controls; whereas alcohol users’ ability is diminished, the effects of cocaine left this ability intact in the current study.

Previous P3AR studies have not reported on the shared variance between theta and delta, and that we found separable results for cocaine users highlights the significance of shared and unique sensitivities in different substances. Our time domain and delta findings showed evidence of a shared process that supports P3AR as a biomarker for broad substance use, as processing was disrupted across both. TF analyses showed that for alcohol users, theta and delta shared variance show a shared process driving activity, whereas cocaine users have unique theta and delta activity, so the effects are driven by different processes. These findings address a need for literature to focus heavily on numerous substances and not predominantly alcohol, to adequately capture and disentangle the unique nature in which substances can affect cognitive processing.

### Differences in P3AR for Target and Novel

It is important to consider that our observed delta effects are stronger in Target trial conditions relative to Novels. Generally, P3 response to Target trials is understood to be reflective of attention and context updating (Polich, 2007), while response to novel stimuli reflects processing of unique versus familiar stimuli (Friedman et al., 2001, 2001; Gaeta et al., 2003; He et al., 2001; Polich, 2007; Spencer et al., 2001; Yago et al., 2003). However, research on the P3 has indicated that Novel and Target P3 responses contain similar, separable frontocentral and centroparietal subcomponents, with the Target P3 evidencing greater activity in the centroparietal subcomponent and the Novel P3 greater in the frontocentral (Dien et al., 2003). Previous research has evaluated ERP activity to novel and target stimuli using time–frequency methods, and findings suggest that the anterior to posterior processing sequence also varies in frequency, with both delta and theta activities having been related to oddball target responses (Bachman & Bernat, 2018; Başar-Eroglu et al., 1992, 2001; Başar-Eroglu & Demiralp, 2001).

This midfrontal theta activity generally precedes delta, and reflects attentional orienting, whereas centroparietal delta activity is closely tied to cognitive processing. Thus, the current study’s observation of greater between-group centroparietal delta effects in Targets relative to Novels is consistent with greater delta activity underlying P3 relative to theta, driving the amplitude reductions (Bachman & Bernat, 2018; Güntekin & Başar, 2016). As delta is comprised mostly of the P3 (Polich, 2007; Basar-Eroglu et al., 1992; Basar-Eroglu et al., 2001; Bernat et al., 2007), it is not surprising that we observed broad P3AR in both the time domain and delta band across substance using groups. In contrast, theta is the frequency band most selectively enhanced for novel stimuli (Cavanagh et al., 2012; Cavanagh & Frank, 2014; Clayton et al., 2015; Karakaş, 2020), thus our weaker novel effects effects are consistent with the contrasting theta responses between Alcohol and Cocaine groups: theta was not diminished for the Cocaine group, but showed reductions for Alcohol group.

Considering these effects in conjunction with the unique physiological effects of alcohol and cocaine, stronger findings in Targets may be related to attentional allocation and task demand. Task-relevant Target trials generally require more active and attentive engagement (Duncan-Johnson & Donchin, 1982; Kok, 2001), thus relative to the salient but task-irrelevant Novel condition, cocaine users were more readily able to process stimuli and respond to task demands, whereas the alcohol users had deficits in their ability to allocate attention effectively. These findings highlight the separable and unique processes underlying cocaine and alcohol use, respectively, and how time-frequency analyses can parse apart these details with more precision than conventional time domain.

### Single versus Poly Users

In our findings, we see P3AR for the Poly group nominally falls between alcohol and cocaine users in the theta band, as seen in mean amplitude values for the SUD groups in Targets and Novels. That the Poly group effects fall between the two single-substance alcohol and cocaine groups further suggests that there are unique substance use processes driving theta for these groups. If there was a shared process for theta across groups, it would be more expectable for combined use (Poly) to show the maximum effect, as opposed to intermediate between the groups. Our findings support the notion that Poly group findings represent a mixture of findings between both groups in opposite directions, cancelling each other out. These findings suggest a blend of mixed effects in both alcohol users with reduced theta and cocaine users with no changes in theta, relative to HC. This supports the view that while delta indexes a shared process of reduced amplitude across cocaine and alcohol use groups, theta represents a unique process that appears to be more modulated for alcohol than cocaine. As there are few ERP studies exploring the P3AR in dually-dependent or concurrently-using subjects, future research should measure more explicitly comorbid substance use to assess the separable impacts of use groups on theta and delta.

### Future Directions

The findings in this paper of a shared P3AR across substances in the time domain, but unique effects for respective substances in time-frequency analyses yield potential for a deeper understanding of shared and unique processes involved in the P3AR as a biomarker for different substances, as opposed to the main focus in the literature on alcohol. Our study found delta and theta activity providing shared and unique sensitivity to cocaine and alcohol use. The study’s findings of separable contributions of theta and delta activity for cocaine and alcohol use, respectively, provide key information that P3 response in other illicit substances may not be modulated in a uniform pattern similar to alcohol, and while there have been conventional time domain findings of P3AR in relation to alcohol, illicit drugs, and nicotine dependence (Patrick et al., 2006), the time frequency analysis focused on unique cocaine and alcohol effects highlight a still-extant dearth of information on interrelationships with the P3AR, thus more ERP studies on a broader array of drugs is clearly necessary.

Further, there are not many investigations of P3AR and substance use utilizing time-frequency analytic approaches. Our findings provide evidence of the importance of TF approaches in separating delta and theta activity to index separable processes underlying P3AR. Thus, future studies would benefit from increased use of time-frequency to separate contributions to P3AR. An additional opportunity when shifting to time-frequency, is that separable activity can also assessed in higher frequencies such as beta, gamma, and alpha frequency bands, which have been well related to ERP activity during the P300 (Almeida et al., 2011; Ford et al., 2008; Polich, 1997; Studenova et al., 2023). Further, alpha, beta, and gamma frequencies have been shown to index relevant processes more broadly, such as relaxation, alertness, and selective attention, respectively, thus indexing processes associated with substance use in these bands is important for future research. There is some recent evidence exploring this relationship, with some work suggesting substance use is associated with increased beta power, for example, but methodological differences and inconsistent findings still render the literature inconclusive (Ceballos et al., 2009; Liu et al., 2022; Ramlakhan et al., 2021). For example, there is little research on cocaine use in these bands, and existing research has demonstrated mixed results (Alper et al., 1990; Bauer, 1994; Herning et al., 1997). For alcohol use, there is much reported in these bands in the resting state, but for the P300 ERP, the relationship within these bands is largely unexplored (Ceballos et al., 2009; Porjesz and Begleiter, 2003). More comprehensive assessment of these processes is important for understanding complex dynamics occurring during P3AR, what processes are modulated, and the degree of shared versus unique variance associated with the processes.

## Conclusion

The present study replicated previous P3AR findings in the traditional time-domain. Most notably, we have extended conventional literature that has focused largely on the association of P3AR with alcohol use to another widely used substance: cocaine. Using time-frequency analyses, we directly compared P3 activity between individuals with single-substance and dually-dependent alcohol and cocaine users, finding broad delta amplitude reductions, and theta amplitude reductions specific to alcohol users. This study specifically examining the P3AR and substance use association with cocaine is novel in the literature, and future work should explore confirmation that differential neurophysiological effects on brain systems account for unique theta sensitivities for cocaine users relative to alcohol. As theta is thought to index attentional orienting, finding unique theta modulations in alcohol users and no-such reductions in cocaine users critically enhances theoretical clarity in understanding P3AR in relation to substance use more broadly. This paper highlights the advantages of time-frequency analytic approaches to more precisely and effectively parse overlapping components otherwise unaccounted for in the traditional time-domain, and advances critical knowledge regarding differential brain system modulations respective to different substances.

## References

Alcohol use. (n.d.). Retrieved February 21, 2024, from https://www.healthdata.org/research-analysis/health-risks-issues/alcohol-use

Almeida, P. R., Vieira, J. B., Silveira, C., Ferreira-Santos, F., Chaves, P. L., Barbosa, F., & Marques-Teixeira, J. (2011). Exploring the dynamics of P300 amplitude in patients with schizophrenia. International Journal of Psychophysiology, PROCEEDINGS OF THE 15TH WORLD CONGRESS OF PSYCHOPHYSIOLOGY of the International Organization of Psychophysiology (I.O.P.) Budapest, Hungary *September 1-4,* 2010, 81(3), 159–168. 10.1016/j.ijpsycho.2011.06.006

Alper, K. R. (1999). The EEG and cocaine sensitization: A hypothesis. The Journal of Neuropsychiatry and Clinical Neurosciences, 11(2), 209–221. 10.1176/jnp.11.2.209

Alper, K. R., Chabot, R. J., Kim, A. H., Prichep, L. S., & John, E. R. (1990). Quantitative EEG correlates of crack cocaine dependence. Psychiatry Research: Neuroimaging, 35(2), 95–105. 10.1016/0925-4927(90)90013-V

Anderson, N. E., Steele, V. R., Maurer, J. M., Bernat, E. M., & Kiehl, K. A. (2015). Psychopathy, attention, and oddball target detection: New insights from PCL-R facet scores. Psychophysiology, 52(9), 1194–1204. 10.1111/psyp.12441

Andrew, C., & Fein, G. (2010). Event-related oscillations vs. Event-related potentials in a P300 task as biomarkers for alcoholism. Alcoholism, Clinical and Experimental Research, 34(4), 669–680. 10.1111/j.1530-0277.2009.01136.x

Anokhin, A. P., Vedeniapin, A. B., Sirevaag, E. J., Bauer, L. O., O’Connor, S. J., Kuperman, S., Porjesz, B., Reich, T., Begleiter, H., Polich, J., & Rohrbaugh, J. W. (2000). The P300 brain potential is reduced in smokers. Psychopharmacology, 149(4), 409–413. 10.1007/s002130000387

Bachman, M. D., & Bernat, E. M. (2018). Independent contributions of theta and delta time-frequency activity to the visual oddball P3b. International Journal of Psychophysiology, 128, 70–80. 10.1016/j.ijpsycho.2018.03.010

Başar-Eroglu, C., Başar, E., Demiralp, T., & Schürmann, M. (1992). P300-response: Possible psychophysiological correlates in delta and theta frequency channels. A review. International Journal of Psychophysiology: Official Journal of the International Organization of Psychophysiology, 13(2), 161–179. 10.1016/0167-8760(92)90055-g

Başar-Eroglu, C., & Demiralp, T. (2001). Event-related theta oscillations: An integrative and comparative approach in the human and animal brain. International Journal of Psychophysiology: Official Journal of the International Organization of Psychophysiology, 39(2–3), 167–195. 10.1016/s0167-8760(00)00140-9

Başar-Eroglu, C., Demiralp, T., Schürmann, M., & Başar, E. (2001). Topological distribution of oddball ‘P300’ responses. International Journal of Psychophysiology, 39(2), 213–220. 10.1016/S0167-8760(00)00142-2

Bashore, T. R., & van der Molen, M. W. (1991). Discovery of the P300: A tribute. Biological Psychology, 32(2–3), 155–171. 10.1016/0301-0511(91)90007-4

Bauer, L. O. (1994). Vigilance in recovering cocaine-dependent and alcohol-dependent patients: A prospective study. Addictive Behaviors, 19(6), 599–607. 10.1016/0306-4603(94)90015-9

Bauer, L. O. (1997). Frontal P300 decrements, childhood conduct disorder, family history, and the prediction of relapse among abstinent cocaine abusers. Drug and Alcohol Dependence, 44(1), 1–10. 10.1016/S0376-8716(96)01311-7

Bauer, L. O. (2001). CNS recovery from cocaine, cocaine and alcohol, or opioid dependence: A P300 study. Clinical Neurophysiology, 112(8), 1508–1515. 10.1016/S1388-2457(01)00583-1

Bauer, L. O. (2002). Differential effects of alcohol, cocaine, and opioid abuse on event-related potentials recorded during a response competition task. Drug and Alcohol Dependence, 66(2), 137–145. 10.1016/S0376-8716(01)00190-9

Bauer, L. O., Costa, L., & Hesselbrock, V. M. (2001). Effects of alcoholism, anxiety and depression on P300 in women: A pilot study. Journal of Studies on Alcohol, 62(5), 571–579. 10.15288/jsa.2001.62.571

Bauer, L. O., & Hesselbrock, V. M. (1999). Subtypes of family history and conduct disorder: Effects on P300 during the stroop test. Neuropsychopharmacology: Official Publication of the American College of Neuropsychopharmacology, 21(1), 51–62. 10.1016/S0893-133X(98)00139-0

Begleiter, H., Porjesz, B., Bihari, B., & Kissin, B. (1984). Event-Related Brain Potentials in Boys at Risk for Alcoholism. Science, 225(4669), 1493–1496. 10.1126/science.6474187

Bernat, E. M., Ellis, J. S., Bachman, M. D., & Hicks, B. M. (2020). P3 Amplitude Reductions are Associated with Shared Variance Between Internalizing and Externalizing Psychopathology. Psychophysiology, 57(7), e13618. 10.1111/psyp.13618

Bernat, E. M., Malone, S. M., Williams, W. J., Patrick, C. J., & Iacono, W. G. (2006). Decomposing delta, theta, and alpha time–frequency ERP activity from a visual oddball task using PCA. International Journal of Psychophysiology : Official Journal of the International Organization of Psychophysiology, 64(1), 62. 10.1016/j.ijpsycho.2006.07.015

Bernat, E. M., Malone, S. M., Williams, W. J., Patrick, C. J., & Iacono, W. G. (2007). Decomposing delta, theta, and alpha time–frequency ERP activity from a visual oddball task using PCA. International Journal of Psychophysiology : Official Journal of the International Organization of Psychophysiology, 64(1), 62–74. 10.1016/j.ijpsycho.2006.07.015

Bernat, E. M., Nelson, L. D., & Baskin-Sommers, A. R. (2015). Time-Frequency Theta and Delta Measures Index Separable Components of Feedback Processing in a Gambling Task. Psychophysiology, 52(5), 626–637. 10.1111/psyp.12390

Bernat, E. M., Nelson, L. D., Steele, V. R., Gehring, W. J., & Patrick, C. J. (2011). Externalizing Psychopathology and Gain/Loss Feedback in a Simulated Gambling Task: Dissociable Components of Brain Response Revealed by Time-Frequency Analysis. Journal of Abnormal Psychology, 120(2), 352–364. 10.1037/a0022124

Bernat, E. M., Williams, W. J., & Gehring, W. J. (2005). Decomposing ERP time–frequency energy using PCA. Clinical Neurophysiology, 116(6), 1314–1334. 10.1016/j.clinph.2005.01.019

Biggins, C. A., MacKay, S., Clark, W., & Fein, G. (1997). Event-related potential evidence for frontal cortex effects of chronic cocaine dependence. Biological Psychiatry, 42(6), 472–485. 10.1016/S0006-3223(96)00425-8

Brennan, G. M., & Baskin-Sommers, A. R. (2018). Brain-behavior relationships in externalizing: P3 amplitude reduction reflects deficient inhibitory control. Behavioural Brain Research, 337, 70–79. 10.1016/j.bbr.2017.09.045

Cavanagh, J. F., & Frank, M. J. (2014). Frontal theta as a mechanism for cognitive control. Trends in Cognitive Sciences, 18(8), 414–421. 10.1016/j.tics.2014.04.012

Cavanagh, J. F., Zambrano-Vazquez, L., & Allen, J. J. B. (2012). Theta lingua franca: A common mid-frontal substrate for action monitoring processes. Psychophysiology, 49(2), 220–238. 10.1111/j.1469-8986.2011.01293.x

Ceballos, N. A., Bauer, L. O., & Houston, R. J. (2009). Recent EEG and ERP Findings in Substance Abusers. Clinical EEG and Neuroscience : Official Journal of the EEG and Clinical Neuroscience Society (ENCS), 40(2), 122. 10.1177/155005940904000210

Ciucă Anghel, D.-M., Nițescu, G. V., Tiron, A.-T., Guțu, C. M., & Baconi, D. L. (2023). Understanding the Mechanisms of Action and Effects of Drugs of Abuse. Molecules, 28(13), 4969. 10.3390/molecules28134969

Clayton, M. S., Yeung, N., & Kadosh, R. C. (2015). The roles of cortical oscillations in sustained attention. Trends in Cognitive Sciences, 19(4), 188–195. 10.1016/j.tics.2015.02.004

Cohen, H. L., Ji, J., Chorlian, D. B., Begleiter, H., & Porjesz, B. (2002). Alcohol-Related ERP Changes Recorded From Different Modalities: A Topographic Analysis. Alcoholism: Clinical and Experimental Research, 26(3), 303–317. 10.1111/j.1530-0277.2002.tb02539.x

Cohen, H. L., Wang, W., Porjesz, B., & Begleiter, H. (1995). Auditory P300 in Young Alcoholics: Regional Response Characteristics. Alcoholism: Clinical and Experimental Research, 19(2), 469–475. 10.1111/j.1530-0277.1995.tb01533.x

Comerchero, M. D., & Polich, J. (1999). P3a and P3b from typical auditory and visual stimuli. Clinical Neurophysiology, 110(1), 24–30. 10.1016/S0168-5597(98)00033-1

Delorme, A., & Makeig, S. (2004). EEGLAB: An open source toolbox for analysis of single-trial EEG dynamics including independent component analysis. Journal of Neuroscience Methods, 134(1), 9–21. 10.1016/j.jneumeth.2003.10.009

Demiralp, T., Ademoglu, A., Istefanopulos, Y., Başar-Eroglu, C., & Başar, E. (2001). Wavelet analysis of oddball P300. International Journal of Psychophysiology: Official Journal of the International Organization of Psychophysiology, 39(2–3), 221–227. 10.1016/s0167-8760(00)00143-4

Di Chiara, G. (1997). Alcohol and Dopamine. Alcohol Health and Research World, 21(2), 108–114.

Dien, J., Spencer, K. M., & Donchin, E. (2003). Localization of the event-related potential novelty response as defined by principal components analysis. Cognitive Brain Research, 17(3), 637–650. 10.1016/S0926-6410(03)00188-5

Diner, B. C., Holcomb, P. J., & Dykman, R. A. (1985). P300 in major depressive disorder. Psychiatry Research, 15(3), 175–184. 10.1016/0165-1781(85)90074-5

Division (DCD), D. C. (2022, November 15). Opioid Facts and Statistics [Text]. https://www.hhs.gov/opioids/statistics/index.html

Duncan-Johnson, C. C., & Donchin, E. (1982). The P300 component of the event-related brain potential as an index of information processing. Biological Psychology, 14(1), 1–52. 10.1016/0301-0511(82)90016-3

Fairbairn, C. E., Kang, D., & Federmeier, K. D. (2021). Alcohol and Neural Dynamics: A Meta-analysis of Acute Alcohol Effects on Event-Related Brain Potentials. *Biological Psychiatry*, Addiction: Reward Learning and Neuroplasticity, 89(10), 990–1000. 10.1016/j.biopsych.2020.11.024

Fillmore, M. T., Rush, C. R., & Hays, L. (2005). Cocaine improves inhibitory control in a human model of response conflict. Experimental and Clinical Psychopharmacology, 13(4), 327–335. 10.1037/1064-1297.13.4.327

Fillmore, M. T., Rush, C. R., & Hays, L. (2006). Acute effects of cocaine in two models of inhibitory control: Implications of non-linear dose effects. Addiction, 101(9), 1323–1332. 10.1111/j.1360-0443.2006.01522.x

First, M. B., Williams, J. B. W., Karg, R. S., & Spitzer, R. L. (2016). *User’s guide for the SCID-5-CV Structured Clinical Interview for DSM-5® disorders: Clinical version* (pp. xii, 158). American Psychiatric Publishing, Inc.

Ford, J. M., Roach, B. J., Hoffman, R. S., & Mathalon, D. H. (2008). The dependence of P300 amplitude on gamma synchrony breaks down in schizophrenia. Brain Research, 1235, 133–142. 10.1016/j.brainres.2008.06.048

Franken, I. H. A., Dietvorst, R. C., Hesselmans, M., Franzek, E. J., Van De Wetering, B. J. M., & Van Strien, J. W. (2008). CLINICAL STUDY: Cocaine craving is associated with electrophysiological brain responses to cocaine-related stimuli. Addiction Biology, 13(3–4), 386–392. 10.1111/j.1369-1600.2008.00100.x

Friedman, D., Cycowicz, Y. M., & Gaeta, H. (2001). The novelty P3: An event-related brain potential (ERP) sign of the brain’s evaluation of novelty. Neuroscience and Biobehavioral Reviews, 25(4), 355–373. 10.1016/s0149-7634(01)00019-7

Gaeta, H., Friedman, D., & Hunt, G. (2003). Stimulus characteristics and task category dissociate the anterior and posterior aspects of the novelty P3. Psychophysiology, 40(2), 198–208. 10.1111/1469-8986.00022

Gao, Y., & Raine, A. (2009). P3 event-related potential impairments in antisocial and psychopathic individuals: A meta-analysis. Biological Psychology, 82(3), 199–210. 10.1016/j.biopsycho.2009.06.006

Garavan, H., Kaufman, J. N., & Hester, R. (2008). Acute effects of cocaine on the neurobiology of cognitive control. Philosophical Transactions of the Royal Society of London. Series B, Biological Sciences, 363(1507), 3267–3276. 10.1098/rstb.2008.0106

Gilmore, C. S., Malone, S. M., Bernat, E. M., & Iacono, W. G. (2010). Relationship between the P3 Event-Related Potential, Its Associated Time-Frequency Components, and Externalizing Psychopathology. Psychophysiology, 47(1), 123–132. 10.1111/j.1469-8986.2009.00876.x

Glenn, S. W., Parsons, O. A., & Smith, L. T. (1996). ERP responses to target and nontarget visual stimuli in alcoholics from VA and community treatment programs. *Alcohol (Fayetteville*, N.Y.), 13(1), 85–92. 10.1016/0741-8329(95)02018-7

Goldstein, R. Z., Leskovjan, A. C., Hoff, A. L., Hitzemann, R., Bashan, F., Khalsa, S. S., Wang, G.-J., Fowler, J. S., & Volkow, N. D. (2004). Severity of neuropsychological impairment in cocaine and alcohol addiction: Association with metabolism in the prefrontal cortex. Neuropsychologia, 42(11), 1447–1458. 10.1016/j.neuropsychologia.2004.04.002

Güntekin, B., & Başar, E. (2016). Review of evoked and event-related delta responses in the human brain. *International Journal of Psychophysiology*, Research on Brain Oscillations and Connectivity in A New Take-Off State, 103, 43–52. 10.1016/j.ijpsycho.2015.02.001

Hajcak, G., MacNamara, A., & Olvet, D. M. (2010). Event-Related Potentials, Emotion, and Emotion Regulation: An Integrative Review. Developmental Neuropsychology, 35(2), 129–155. 10.1080/87565640903526504

Hamidovic, A., & Wang, Y. (2019). The P300 in alcohol use disorder: A meta-analysis and meta-regression. Progress in Neuro-Psychopharmacology and Biological Psychiatry, 95, 109716. 10.1016/j.pnpbp.2019.109716

Harper, J., Malone, S. M., Bachman, M. D., & Bernat, E. M. (2016). Stimulus sequence context differentially modulates inhibition-related theta and delta band activity in a go/nogo task. Psychophysiology, 53(5), 712. 10.1111/psyp.12604

Harper, J., Malone, S. M., & Bernat, E. M. (2014). Theta and delta band activity explain N2 and P3 ERP component activity in a go/no-go task. Clinical Neurophysiology, 125(1), 124–132. 10.1016/j.clinph.2013.06.025

He, B., Lian, J., Spencer, K. M., Dien, J., & Donchin, E. (2001). A cortical potential imaging analysis of the P300 and Novelty P3 components. Human Brain Mapping, 12(2), 120–130. 10.1002/1097-0193(200102)12:2%253C120::AID-HBM1009%253E3.0.CO;2-V

Herning, R. I., King, D. E., Better, W., & Cadet, J. L. (1997). Cocaine Dependence. Annals of the New York Academy of Sciences, 825(1), 323–327. 10.1111/j.1749-6632.1997.tb48442.x

Hesselbrock, V., Begleiter, H., Porjesz, B., O’Connor, S., & Bauer, L. (2001). P300 event-related potential amplitude as an endophenotype of alcoholism—Evidence from the collaborative study on the genetics of alcoholism. Journal of Biomedical Science, 8(1), 77–82. 10.1007/BF02255974

Hosch, A., Harris, J. L., Swanson, B., & Petersen, I. T. (2023). The P3 ERP in Relation to General Versus Specific Psychopathology in Early Childhood. Research on Child and Adolescent Psychopathology, 51(10), 1439–1451. 10.1007/s10802-023-01061-0

Iacono, W. G., Carlson, S. R., Malone, S. M., & McGue, M. (2002). P3 event-related potential amplitude and the risk for disinhibitory disorders in adolescent boys. Archives of General Psychiatry, 59(8), 750–757. 10.1001/archpsyc.59.8.750

Iacono, W. G., Malone, S. M., & McGue, M. (2003). Substance use disorders, externalizing psychopathology, and P300 event-related potential amplitude. International Journal of Psychophysiology: Official Journal of the International Organization of Psychophysiology, 48(2), 147–178. 10.1016/s0167-8760(03)00052-7

Jasper, H.H. (1958). The Ten-Twenty Electrode System of the International Federation. Electroencephalography and Clinical Neurophysiology, 10, 371–375.

Jeon, Y.-W., & Polich, J. (2001). P300 asymmetry in schizophrenia: A meta-analysis. Psychiatry Research, 104(1), 61–74. 10.1016/S0165-1781(01)00297-9

Jeon, Y.-W., & Polich, J. (2003). Meta-analysis of P300 and schizophrenia: Patients, paradigms, and practical implications. Psychophysiology, 40(5), 684–701. 10.1111/1469-8986.00070

Jones, K. A., Porjesz, B., Chorlian, D., Rangaswamy, M., Kamarajan, C., Padmanabhapillai, A., Stimus, A., & Begleiter, H. (2006). S-transform time-frequency analysis of P300 reveals deficits in individuals diagnosed with alcoholism. Clinical Neurophysiology, 117(10), 2128–2143. 10.1016/j.clinph.2006.02.028

Jung, T. P., Makeig, S., Humphries, C., Lee, T. W., McKeown, M. J., Iragui, V., & Sejnowski, T. J. (2000). Removing electroencephalographic artifacts by blind source separation. Psychophysiology, 37(2), 163–178.

Karakaş, S. (2020). A review of theta oscillation and its functional correlates. International Journal of Psychophysiology, 157, 82–99. 10.1016/j.ijpsycho.2020.04.008

Karakaş, S., Erzengin, Ö. U., & Başar, E. (2000). A new strategy involving multiple cognitive paradigms demonstrates that ERP components are determined by the superposition of oscillatory responses. Clinical Neurophysiology, 111(10), 1719–1732. 10.1016/S1388-2457(00)00418-1

Keil, A., Bernat, E. M., Cohen, M. X., Ding, M., Fabiani, M., Gratton, G., Kappenman, E. S., Maris, E., Mathewson, K. E., Ward, R. T., & Weisz, N. (2022). Recommendations and publication guidelines for studies using frequency domain and time-frequency domain analyses of neural time series. Psychophysiology, 59(5), e14052. 10.1111/psyp.14052

Kempel, P., Lampe, K., Parnefjord, R., Hennig, J., & Kunert, H. J. (2003). Auditory-Evoked Potentials and Selective Attention: Different Ways of Information Processing in Cannabis Users and Controls. Neuropsychobiology, 48(2), 95–101. 10.1159/000072884

Kessler, R. C., Crum, R. M., Warner, L. A., Nelson, C. B., Schulenberg, J., & Anthony, J. C. (1997). Lifetime co-occurrence of DSM-III-R alcohol abuse and dependence with other psychiatric disorders in the National Comorbidity Survey. Archives of General Psychiatry, 54(4), 313–321. 10.1001/archpsyc.1997.01830160031005

Kiehl, K. A., Laurens, K. R., Duty, T. L., Forster, B. B., & Liddle, P. F. (2001). Neural sources involved in auditory target detection and novelty processing: An event-related fMRI study. Psychophysiology, 38(1), 133–142.

Kimble, M., Kaloupek, D., Kaufman, M., & Deldin, P. (2000). Stimulus novelty differentially affects attentional allocation in PTSD. Biological Psychiatry, 47(10), 880–890. 10.1016/s0006-3223(99)00258-9

Klimesch, W. (1999). EEG alpha and theta oscillations reflect cognitive and memory performance: A review and analysis. Brain Research. Brain Research Reviews, 29(2–3), 169–195. 10.1016/s0165-0173(98)00056-3

Kok, A. (2001). On the utility of P3 amplitude as a measure of processing capacity. Psychophysiology, 38(3), 557–577. 10.1017/S0048577201990559

Kotov, R., Krueger, R. F., Watson, D., Achenbach, T. M., Althoff, R. R., Bagby, R. M., Brown, T. A., Carpenter, W. T., Caspi, A., Clark, L. A., Eaton, N. R., Forbes, M. K., Forbush, K. T., Goldberg, D., Hasin, D., Hyman, S. E., Ivanova, M. Y., Lynam, D. R., Markon, K., … Zimmerman, M. (2017). The Hierarchical Taxonomy of Psychopathology (HiTOP): A dimensional alternative to traditional nosologies. Journal of Abnormal Psychology, 126(4), 454–477. 10.1037/abn0000258

Kouri, E. M., Lukas, S. E., & Mendelson, J. H. (1996). P300 assessment of opiate and cocaine users: Effects of detoxification and buprenorphine treatment. Biological Psychiatry, 40(7), 617–628. 10.1016/0006-3223(95)00468-8

Liu, Y., Chen, Y., Fraga-González, G., Szpak, V., Laverman, J., Wiers, R. W., & Richard Ridderinkhof, K. (2022). Resting-state EEG, Substance use and Abstinence After Chronic use: A Systematic Review. Clinical EEG and Neuroscience, 53(4), 344–366. 10.1177/15500594221076347

Ma, H., & Zhu, G. (2014). The dopamine system and alcohol dependence. Shanghai Archives of Psychiatry, 26(2), 61–68. 10.3969/j.issn.1002-0829.2014.02.002

Mash, D. C., Pablo, J., Ouyang, Q., Hearn, W. L., & Izenwasser, S. (2002). Dopamine transport function is elevated in cocaine users. Journal of Neurochemistry, 81(2), 292–300. 10.1046/j.1471-4159.2002.00820.x

Milandu, J., Pavuluri, A., Mannava, P., Blaise, L., Risco, C., Butler, D., Mello, G., Fix, S., & Bernat, E. (2023). Brain Measures of Shared and Unique Variance between Internalizing and Externalizing Psychopathology. In Program No. PSTR229.07. Neuroscience Meeting.

Molnár, M. (1994). On the origin of the P3 event-related potential component. International Journal of Psychophysiology, 17(2), 129–144. 10.1016/0167-8760(94)90028-0

Nelson, L. D., Patrick, C. J., Collins, P., Lang, A. R., & Bernat, E. M. (2011). Alcohol impairs brain reactivity to explicit loss feedback. Psychopharmacology, 218(2), 419–428. 10.1007/s00213-011-2323-3

Pandey, A. K., Kamarajan, C., Manz, N., Chorlian, D. B., Stimus, A., & Porjesz, B. (2016). Delta, theta, and alpha event-related oscillations in alcoholics during Go/NoGo task: Neurocognitive deficits in execution, inhibition, and attention processing. Progress in Neuro-Psychopharmacology and Biological Psychiatry, 65, 158–171. 10.1016/j.pnpbp.2015.10.002

Patrick, C. J., Bernat, E. M., Malone, S. M., Iacono, W. G., Krueger, R. F., & McGue, M. (2006). P300 amplitude as an indicator of externalizing in adolescent males. Psychophysiology, 43(1), 84–92. 10.1111/j.1469-8986.2006.00376.x

Pennings, E. J. M., Leccese, A. P., & Wolff, F. A. de. (2002). Effects of concurrent use of alcohol and cocaine. Addiction, 97(7), 773–783. 10.1046/j.1360-0443.2002.00158.x

Pike, E., Marks, K. R., Stoops, W. W., & Rush, C. R. (2015). Cocaine-related stimuli impair inhibitory control in cocaine users following short stimulus onset asynchronies. Addiction, 110(8), 1281–1286. 10.1111/add.12947

Polich, J. (1997). On the relationship between EEG and P300: Individual differences, aging, and ultradian rhythms. International Journal of Psychophysiology, 26(1), 299–317. 10.1016/S0167-8760(97)00772-1

Polich, J. (2004). Clinical application of the P300 event-related brain potential. Physical Medicine and Rehabilitation Clinics of North America, 15(1), 133–161. 10.1016/s1047-9651(03)00109-8

Polich, J. (2007). Updating P300: An Integrative Theory of P3a and P3b. Clinical Neurophysiology : Official Journal of the International Federation of Clinical Neurophysiology, 118(10), 2128–2148. 10.1016/j.clinph.2007.04.019

Polich, J., Pollock, V. E., & Bloom, F. E. (1994). Meta-analysis of P300 amplitude from males at risk for alcoholism. Psychological Bulletin, 115(1), 55–73. 10.1037/0033-2909.115.1.55

Porjesz, B., & Begleiter, H. (1995). Event-Related Potentials and Cognitive Function in Alcoholism. Alcohol Health and Research World, 19(2), 108–112.

Porjesz, B., & Begleiter, H. (2003). Alcoholism and Human Electrophysiology. Alcohol Research & Health, 27(2), 153.

Porjesz, B., Begleiter, H., & Garozzo, R. (1980). Visual Evoked Potential Correlates of Information Processing Deficits in Chronic Alcoholics. In Henri Begleiter (Ed.), Biological Effects of Alcohol (pp. 603–623). Springer US. 10.1007/978-1-4684-3632-7_46

Porjesz, B., Begleiter, H., & Samuelly, I. (1980). Cognitive deficits in chronic alcoholics and elderly subjects assessed by evoked brain potentials. Acta Psychiatrica Scandinavica. Supplementum, 286, 15–29. 10.1111/j.1600-0447.1980.tb08051.x

Porjesz, B., Rangaswamy, M., Kamarajan, C., Jones, K. A., Padmanabhapillai, A., & Begleiter, H. (2005). The utility of neurophysiological markers in the study of alcoholism. Clinical Neurophysiology, 116(5), 993–1018. 10.1016/j.clinph.2004.12.016

Ramlakhan, J. U., Ma, M., Zomorrodi, R., Blumberger, D. M., Noda, Y., & Barr, M. S. (2021). The Role of Gamma Oscillations in the Pathophysiology of Substance Use Disorders. Journal of Personalized Medicine, 11(1), Article 1. 10.3390/jpm11010017

Rangaswamy, M., Jones, K. A., Porjesz, B., Chorlian, D. B., Padmanabhapillai, A., Kamarajan, C., Kuperman, S., Rohrbaugh, J., O’Connor, S. J., Bauer, L. O., Schuckit, M. A., & Begleiter., H. (2007). Delta and Theta oscillations as risk markers in Adolescent Offspring of Alcoholics. International Journal of Psychophysiology : Official Journal of the International Organization of Psychophysiology, 63(1), 3–15. 10.1016/j.ijpsycho.2006.10.00

Rangaswamy, M., & Porjesz, B. (2014). Understanding alcohol use disorders with neuroelectrophysiology. Handbook of Clinical Neurology, 125, 383–414. 10.1016/B978-0-444-62619-6.00023-9

Robins, L. N., & Regier, D. A. (1991). *Psychiatric disorders in America: The epidemiologic catchment area study*. Free Press.

Rodriguez Holguin, S., Porjesz, B., Chorlian, D. B., Polich, J., & Begleiter, H. (1999). Visual P3a in male alcoholics and controls. Alcoholism, Clinical and Experimental Research, 23(4), 582–591.

Sara, G., Gordon, E., Kraiuhin, C., Coyle, S., Howson, A., & Meares, R. (1994). The P300 ERP component: An index of cognitive dysfunction in depression? Journal of Affective Disorders, 31(1), 29–38. 10.1016/0165-0327(94)90124-4

Sarafino, E. P., & Smith, T. W. (2022). Health Psychology: Biopsychosocial Interactions. John Wiley & Sons.

Shelton, M. D., & Parsons, O. A. (1987). Alcoholics’ self-assessment of their neuropsychological functioning in everyday life. Journal of Clinical Psychology, 43(3), 395–403. 10.1002/1097-4679(198705)43:3%253C395::AID-JCLP2270430314%253E3.0.CO;2-Z

Sher, K. J., & Trull, T. J. (1994). Personality and disinhibitory psychopathology: Alcoholism and antisocial personality disorder. Journal of Abnormal Psychology, 103(1), 92–102. 10.1037/0021-843X.103.1.92

Siegel, R. K. (1984a). Cocaine and the Privileged Class: A Review of Historical and Contemporary Images. Advances in Alcohol & Substance Abuse, 4(2), 37–49. 10.1300/J251v04n02_04

Siegel, R. K. (1984b). Cocaine smoking disorders: Diagnosis and treatment. Psychiatric Annals, 14(10), 728–732. 10.3928/0048-5713-19841001-08

Singh, R., Shukla, R., Dalal, P. K., Sinha, P. K., & Trivedi, J. K. (2000). P 300 EVENT RELATED POTENTIAL IN DEPRESSION. Indian Journal of Psychiatry, 42(4), 402–409.

Singh, S. M., & Basu, D. (2009). CLINICAL STUDY: The P300 event-related potential and its possible role as an endophenotype for studying substance use disorders: a review. Addiction Biology, 14(3), 298–309. 10.1111/j.1369-1600.2008.00124.x

Singh, S. M., Basu, D., Kohli, A., & Prabhakar, S. (2009). Auditory P300 Event-Related Potentials and Neurocognitive Functions in Opioid Dependent Men and Their Brothers. American Journal on Addictions, 18(3), 198–205. 10.1080/10550490902786975

Spencer, K. M., Dien, J., & Donchin, E. (2001). Spatiotemporal analysis of the late ERP responses to deviant stimuli. Psychophysiology, 38(2), 343–358. 10.1111/1469-8986.3820343

Spronk, D. B., De Bruijn, E. R. A., van Wel, J. H. P., Ramaekers, J. G., & Verkes, R. J. (2016). Acute effects of cocaine and cannabis on response inhibition in humans: An ERP investigation. Addiction Biology, 21(6), 1186–1198. 10.1111/adb.12274

Steele, V. R., Claus, E. D., Aharoni, E., Vincent, G. M., Calhoun, V. D., & Kiehl, K. A. (2015). Multimodal imaging measures predict rearrest. Frontiers in Human Neuroscience, 9. 10.3389/fnhum.2015.00425

Steele, V. R., Fink, B. C., Maurer, J. M., Arbabshirani, M. R., Wilber, C. H., Jaffe, A. J., Sidz, A., Pearlson, G. D., Calhoun, V. D., Clark, V. P., & Kiehl, K. A. (2014). Brain Potentials Measured During a Go/NoGo Task Predict Completion of Substance Abuse Treatment. *Biological Psychiatry*, Dopamine Deficits as an Addiction Phenotype, 76(1), 75–83. 10.1016/j.biopsych.2013.09.030

Studenova, A., Forster, C., Engemann, D. A., Hensch, T., Sanders, C., Mauche, N., Hegerl, U., Loffler, M., Villringer, A., & Nikulin, V. (2023). Event-related modulation of alpha rhythm explains the auditory P300-evoked response in EEG. eLife, 12, RP88367. 10.7554/eLife.88367

Turetsky, B. I., Cannon, T. D., & Gur, R. E. (2000). P300 subcomponent abnormalities in schizophrenia: III. Deficits In unaffected siblings of schizophrenic probands. Biological Psychiatry, 47(5), 380–390. 10.1016/s0006-3223(99)00290-5

UNODC World Drug Report 2022—World | ReliefWeb. (2022, June 27). https://reliefweb.int/report/world/unodc-world-drug-report-2022

Verleger, R. (1988). Event-related potentials and cognition: A critique of the context updating hypothesis and an alternative interpretation of P3. Behavioral and Brain Sciences, 11(3), 343–356. 10.1017/S0140525X00058015

Wan, L., Baldridge, R. M., Colby, A. M., & Stanford, M. S. (2010). Association of P3 amplitude to treatment completion in substance dependent individuals. Psychiatry Research, 177(1), 223–227. 10.1016/j.psychres.2009.01.033

Watts, A. T. M., Bachman, M. D., & Bernat, E. M. (2017). Expectancy effects in feedback processing are explained primarily by time-frequency delta not theta. Biological Psychology, 129, 242–252. 10.1016/j.biopsycho.2017.08.054

Xu, S., Chai, H., Hu, J., Xu, Y., Chen, W., & Wang, W. (2014). Passive Event-Related Potentials to a Single Tone in Treatment-Resistant Depression, Generalized Anxiety Disorder, and Borderline Personality Disorder Patients. Journal of Clinical Neurophysiology, 31(5), 488. 10.1097/WNP.0000000000000091

Yago, E., Escera, C., Alho, K., Giard, M.-H., & Serra-Grabulosa, J. M. (2003). Spatiotemporal dynamics of the auditory novelty-P3 event-related brain potential. Cognitive Brain Research, 16(3), 383–390. 10.1016/S0926-6410(03)00052-1

Yordanova, J., Kolev, V., Heinrich, H., Banaschewski, T., Woerner, W., & Rothenberger, A. (2000). Gamma band response in children is related to task-stimulus processing. NeuroReport, 11(10), 2325–2330.

